# Social cues of safety can override differences in threat level

**DOI:** 10.1101/2022.02.27.481960

**Authors:** Clara H Ferreira, Mirjam Heinemans, Matheus Farias, Rui Gonçalves, Marta A Moita

## Abstract

Animals in groups integrate social information with that directly-gathered about the environment to guide decisions regarding reproduction, foraging and defense against predatory threats. In the context of predation, usage of social information has acute fitness benefits, aiding the detection of predators, the mounting of concerted defensive responses, or allowing the inference of safety, permitting other beneficial behaviors such as foraging for food. Individual and group defense responses to predatory threats can vary in modality and vigor depending on the perceived threat level. Moreover, predation level has been shown to modulate the use of social cues about foraging sites. Whether and how different threat levels affect the use of social cues to guide defense responses, is currently unknown. We previously showed that *Drosophila melanogaster* display a graded decrease in freezing behavior, triggered by an inescapable visual threat, with increasing group sizes. Crucially, we identified the movement of others as a cue of safety and its cessation a cue of threat and found the group responses to be primarily guided by the safety cues, resulting in a net social buffering effect. Here, we investigated how threat level impacts the use of social cues by exposing flies individually and in groups to two threat imminences using looms of different speeds. We show that freezing responses are stronger to the faster looms regardless of social condition. However, social buffering was stronger for groups exposed to the fast looms, such that the increase in freezing caused by the higher threat was less prominent in flies tested in groups than those tested individually. Through artificial control of behavior, we created different group compositions, titrating the motion cues that were maintained across threat levels. We, found that the same level of safety motion cues had a bigger weight on the flies’ decisions when these were exposed to the higher threat, thus overriding differences in perceived threat levels. These findings shed light on the ‘safety in numbers’ effect, revealing the modulation of the saliency of social safety cues across threat intensities, a possible mechanism to regulate costly defensive responses.

## Introduction

A major benefit of being in a group is the possibility of adding social information to directly-perceived information about the environment to guide behavior. Across the animal kingdom, this social information can be actively transmitted via signals evolved specifically for communication ^1,2^, but also acquired through information-bearing cues of different sensory natures, which animals produce as they engage in their daily activities. Vertebrates use such social cues to procure food ^3^ for example using vision to assess where and how much others are eating ^4^, to choose mates by copying the decisions of others ^5^ based for example on olfactory cues ^6^, and to infer predation threat levels ^7^ for instance by auditory detection of escape ^8^ or freezing (active immobility response aimed at becoming inconspicuous) ^9^. These types of social cue usage are also reported in invertebrates including in *Drosophila melanogaster* ^10,11^, guiding aggregation on food ^12–15^, reproduction related decisions in mating ^16,17^ and oviposition ^18–20^, as well defensive responses^21^, many of which relying at least partially on vision.

The acquisition and exploitation of the information provided by social cues can confer fitness benefits ^22^, particularly in the context of a response to a potential threat: failure to detect a predator can lead to an animal’s immediate demise whereas needless engagement in metabolically costly defense responses ^23^ can negatively impact survival. We study these defense responses to threat, which can be triggered by diverse stimuli ^24^, and are subject to modulation by a variety of factors ^25^ from the sensory modality of the threatening stimulus and its properties, to environmental factors including the social environment. We use visual looming stimuli mimicking an approaching predator to elicit defensive responses, which have been reported for all visual animals tested so far, from invertebrates such as crabs ^26^ and flies ^27,28^, to vertebrates like mice ^29^ and humans ^30^. Using visual threats, whose properties can easily be manipulated in the lab, permits a detailed understanding of how different aspects of a threat affect the crucial deployment of defense responses. For example, a black disk sweeping overhead, mimicking a cruising predator, elicits freezing, while the same black disc expanding on the screen, as if looming towards the mouse, induces escapes ^31^. Similarly, fruit flies have been shown to respond both with escapes and with freezing to repeated, inescapable, sweeping and looming stimuli, where escapes predominate in response to sweeps ^32^ and freezing in response to looms ^27^. In addition, in both zebrafish larvae ^33^ and flies ^34,35^ looming stimuli with lower approach rates evoke slower escapes, while higher approach rates evoke faster responses.

We had previously shown that flies exposed to looming stimuli in groups use social motion cues as both a cue of threat and safety, with freezing in others leading to freezing in a focal fly, and movement of others leading to movement resumption after freezing ^21^. Surprisingly, despite the ease of manipulation of the looming stimulus, and the knowledge that the social environment influences the deployment of defense responses, detailed studies of how different threat levels impact group behavior are still scarce. There are some reports of modulation of group responses with threat, showing that prey species from higher-predation habitats form larger and more cohesive groups than those from lower predation environments ^36–39^, and that different degrees of uniformity in escape formations can be triggered by manipulating the angle of approach of a threat ^40^. At the interplay of foraging and predation there are further examples of modulation by the social environment and reliance on social cues ^22^. For example bumblebees use the presence of conspecifics as a cue of safety, joining others at foraging sites only in potentially hazardous situations, when those sites were previously predator-infested ^41^. Similarly, minnows use socially-derived information and copy feeding location to a higher extent when exposed to a higher predation risk ^42^, an example of the ‘copy-when-asocial-learning-is-costly’ hypothesis regarding social learning. These examples point to the fact that group behaviors are modulated by threat level, and that reliance on social cues also varies with predation risk. However, how different threat levels affect group defensive responses, such as freezing, and the reliance on social cues is unknown.

In this study we use two loom speeds as threat manipulation and show that faster looms induce higher freezing responses both in individually tested flies and flies in groups, but that the group effect on freezing responses depends on threat imminence. We then manipulate the social environment, controlling the numbers of moving and freezing flies surrounding a focal fly to clearly disentangle the effect of looming speed and social motion cue usage on freezing responses. We find that flies display graded, decreasing, freezing responses with increasing numbers of moving flies and hence motion cues, for both looming speeds. Though freezing levels are generally higher for faster looms, differences in freezing responses collapse across looming speeds as soon as at least one surrounding fly is moving. Our findings suggest that social motion cues of safety have a preponderant role in freezing responses, which overrides differences in threat level.

## Materials and methods

### Fly lines and husbandry

Flies were kept at 25 °C and 70% humidity in a 12 h:12 h dark:light cycle. Experimental animals were mated females, tested only once when 4–6 days old. For optogenetic manipulations flies were transferred for 48h before the experiments to food with 0.4 mmol/L retinal. For experiment with mixed genotypes, focal flies were marked on the thorax using white marker pen.

Wild-type flies used were Canton-S. LC6-splitGAL4 line *w[1118]; P{y[+t7.7] w[+mC]=R92B02-p65.AD}attP40; P{y[+t7.7] w[+mC]=R41C07-GAL4.DBD}attP2*^43^ and *w[*] norpA[36]* were obtained from the Bloomington stock center. *UAS-Chrimson* line used was *w*^1118^; *P{20XUAS-IVS-CsChrimson.mVenus}attP2*^44^.

### Behavioral apparatus

Behavioral experiments were performed as described in ^21^, using an updated setup (Farias et al., in preparation). Briefly, we imaged unrestrained flies in 11° slanted PETG arenas with 68 mm diameter (central flat portion diameter 32 mm) while being presented with visual stimulation (twenty 500 ms looming stimuli, a black circle in a white background, exponentially expanding at a speed 25 or 50 cm/s) on an Asus monitor running at 240 Hz, tilted 45° over the stage (Figure 1A). The stage contained two arenas and under it a custom-built LED board that provided blacklight for imaging (infrared, 940nm) and red-light (627nm) for optogenetic stimulation. Videos were acquired through Bonsai ^45^ at 60 Hz and 1280 width × 960 height resolution using two USB3 cameras (PointGrey Flea3).

**Figure 1.**
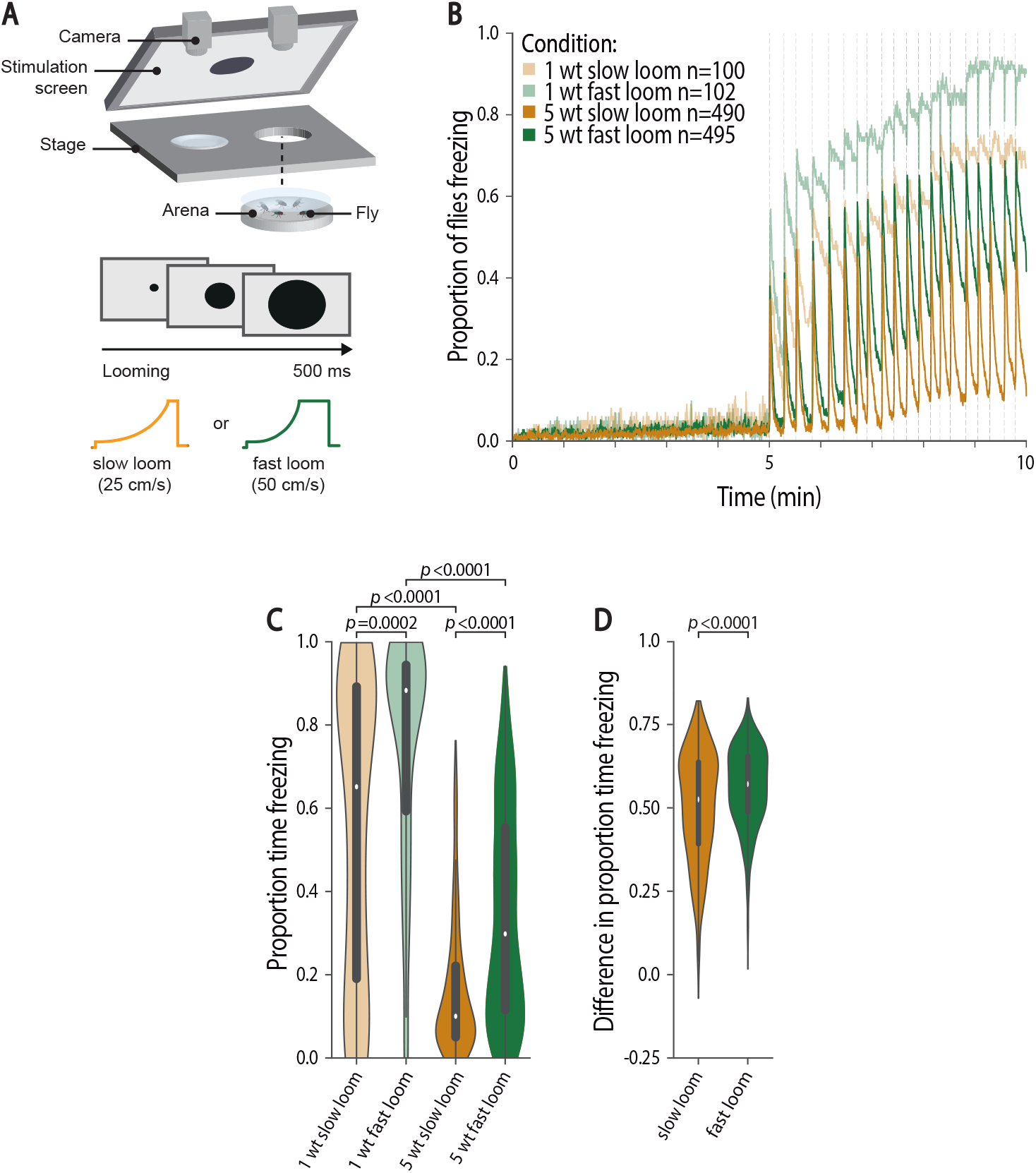
Group freezing responses scale with threat imminence. A) Experimental setup and conditions: we tested individuals and groups of five flies in backlit arenas imaged from above; after a 5-min baseline flies were exposed to twenty 500 ms looming presentations, every 10–20 s; we provided either slow (25 cm/s, purple) or fast looms (50 cm/s, green). B-C) Data for flies tested individually (lighter shades) and in groups (darker shades). B) Fraction of flies freezing throughout the experiment; dashed lines represent looming stimuli presentations. C-D) Violin plots representing the probability density distribution of individual fly data bound to the range of possible values, with boxplots elements: center line, median; box limits, upper (75) and lower (25) quartiles; whiskers, 1.5x interquartile range). C) Proportion of time spent freezing in the stimulation period. P-values result from Kruskal–Wallis statistical analysis followed by Dunn’s multiple comparisons test. D) Difference in the proportion of time spent freezing between individually tested flies and flies tested in groups for slow and fast looms (see Methods). P-value results from two-tailed Mann–Whitney test.

Optogenetic manipulation of LC6>CsChrimson without concurrent loom presentations were done during a 5 min stimulation period, after a 5 min baseline period, with the application of 20 stimuli of pulsed red light at 50Hz, 50% duty cycle (DC), 7.5 mw/cm2 normalized intensity. Optogenetic manipulations with simultaneous looming stimuli were performed during a 2-minute period, at 50Hz, 50% DC, 10 mW/cm^2^ normalized intensity.

### Data analysis

Data were analyzed using custom scripts in spyder (python 3.8). Statistical testing was done in GraphPad Prism 7.03, and non-parametric, Kruskal-Wallis followed by Dunn’s multiple comparison test or two-tailed Mann-Whitney tests were chosen, as data were not normally distributed (Shapiro-Wilk test).

Freezing bouts were classified as zero-pixel change detected in a 4×4 mm square around the fly for at least 500 ms (30 frames). To correct for jitter in the pixel change, a maximum break of 50 ms (3 frames) was allowed within freezing bouts; any pixel change detected for more than this period was considered a break in freezing. Freezing bout onset around loom was determined from 30 frames before each loom until 150 frames after.

The proportion of time spent freezing was quantified by taking the sum of the frames in which freezing occurred during the stimulation period and dividing that by the total number of frames of this period. Freezing bout lengths reflect the freezing duration during the stimulation period, from onset to offset or until the experiment ended. The proportion of freezing exits between looms was calculated by determining whether a fly froze upon the 2 s after looming and assessing its freezing status in the last 0.5 s before the next loom.

To make pairwise comparisons of the proportion of time spent freezing and of freezing exits between conditions (flies tested individually vs. in groups, or flies exposed to slow vs. fast looms) we created 1.000 difference values, where each data point corresponds to the difference between the median of 20 samples, with replacement, of a behavioral measure from each condition (for example: median of 20 samples of time freezing of individually tested flies – median of 20 samples of time spent freezing of group tested flies).

To determine when a jump occurred, we identified when a fly’s speed exceeded 75 mm/s for at least one frame, and applied a time constraint of 3 frames between two consecutive jumps.

### Logistic regression model

We modelled the decision to stay frozen or resume movement using the scikit-learn logistic regression model, as previously described ^21^. Briefly, we modelled the probability of exiting freezing in between looming stimuli as a function of the looming speed and the average of the sum of the motion cue generated by neighboring flies during that freezing bout. As previously described, we calculated motion cues for a focal as the summed product of speed and angle on the retina of a focal fly that each other fly produces *∑ speed* × *anqle on the retina* (*θ*) where 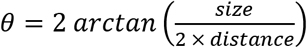. We used 10,0000 times bootstrapped data with replacement. To determine the explanatory power of each predictor, we determined the associated fraction of variance.

## Results

To study how different threat imminences affect flies’ social defensive responses, we first assessed whether the speed with which a looming dark disc approaches modulates freezing responses. To this end, we compared freezing behavior of individually tested flies exposed to one of two looming speeds, 25 and 50 cm/s (Figure 1 A). A pilot experiment in the lab suggested individual freezing responses varied with looming speed and, indeed, we found that the fraction of flies freezing throughout the experiment when exposed to twenty renditions of the slower loom (25 cm/s) is inferior to that observed when flies are exposed to the same number of renditions at a faster speed (50 cm/s) (Figure 1B). This is further corroborated by the increased time spent freezing by flies exposed to the faster looms compared to flies exposed to the slower looms (PropF_25_ = 0.65 IQR 0.18–0.89, Prop_F50_ = 0.88 IQR 0.58–0.94, Kruskal–Wallis, KW, followed by Dunn’s multiple comparisons test, D, p = 0.0002; Figure 1C). In concordance with previous results for escapes ^33^, exposure to faster looms led to more rapid freezing responses, as revealed by the shorter latencies to freeze in response to the fast looms relative to slow looms (Supplementary Figure 1A). Having established that increasing looming speed increases freezing responses, we will henceforth use looming speed to study how different levels of threat affect social defensive behavior.

As flies show social buffering of defensive responses, that is, when exposed to a threat while surrounded by others they freeze less ^21^ and groups of animals can behave differently depending on the level of threat ^36–40^, we hypothesized that experiencing looms of different speeds impacts group behavior and the weights given to the available social information. We thus compared freezing responses of flies, tested in groups of five, exposed to the fast and slow looms (Figure 1B, C) and found that, just as individually tested flies, groups of five flies exposed to the faster looms freeze more than groups exposed to slower looms (Prop_F25_ = 0.10 IQR 0.50–0.22, Prop_F50_ = 0.30 IQR 0.11–0.55, KWD p < 0.0001; Figure 1C). In addition, flies exposed to fast looms in groups show shorter latencies to start freezing that are similar to those observed for flies tested individually (Supplementary Figure 1B). Although both flies tested individually and flies tested in groups responded with stronger freezing to the faster loom, it is still possible that the impact of the social environment, that is the degree of social buffering, varied with threat level. To test this possibility, we compared the decrease in freezing of flies tested in groups relative to freezing levels of individually tested flies for both loom speeds (see Methods, Figure 1D) and found that the decrease in freezing, caused by social buffering, was bigger for flies exposed to fast looms (two-tailed Mann-Whitney, MW, p<0.0001). In summary, the time flies spend freezing scales with perceived threat imminence, but the social environment seems to have a higher weight in guiding freezing responses when flies are exposed to faster looms.

When in a group, the movement generated by the neighboring flies leads to an increase in freezing exits resulting in faster resumption of activity and less sustained freezing in between loom presentations ^21^. Hence, we analyzed the effect of looming speed on the probability of exiting freezing (Figure 2). Consistent with our previous results, flies tested individually display low probability of exiting freezing (Prob(Fexit)_25_ = 0.21, IQR 0.06–0.67, Prob(Fexit)_50_ = 0.08 IQR 0.05–0.17; Figure 2A), and flies tested in groups are more likely to stop freezing in between looming stimuli (Prob(Fexit)_25_ = 0.94 IQR 0.82–1.00, KWD p < 0.0001, Prob(Fexit)_50_ = 0.79 IQR 0.44–0.94, KWD p < 0.0001; Figure 2A). In line with the results for the proportion of time spent freezing (Figure 1), perceived threat imminence significantly affects the probability of exiting freezing in flies tested individually (KWD p = 0.0004; Figure 2A) and in groups (KWD p < 0.0001; Figure 2A).

**Figure 2.**
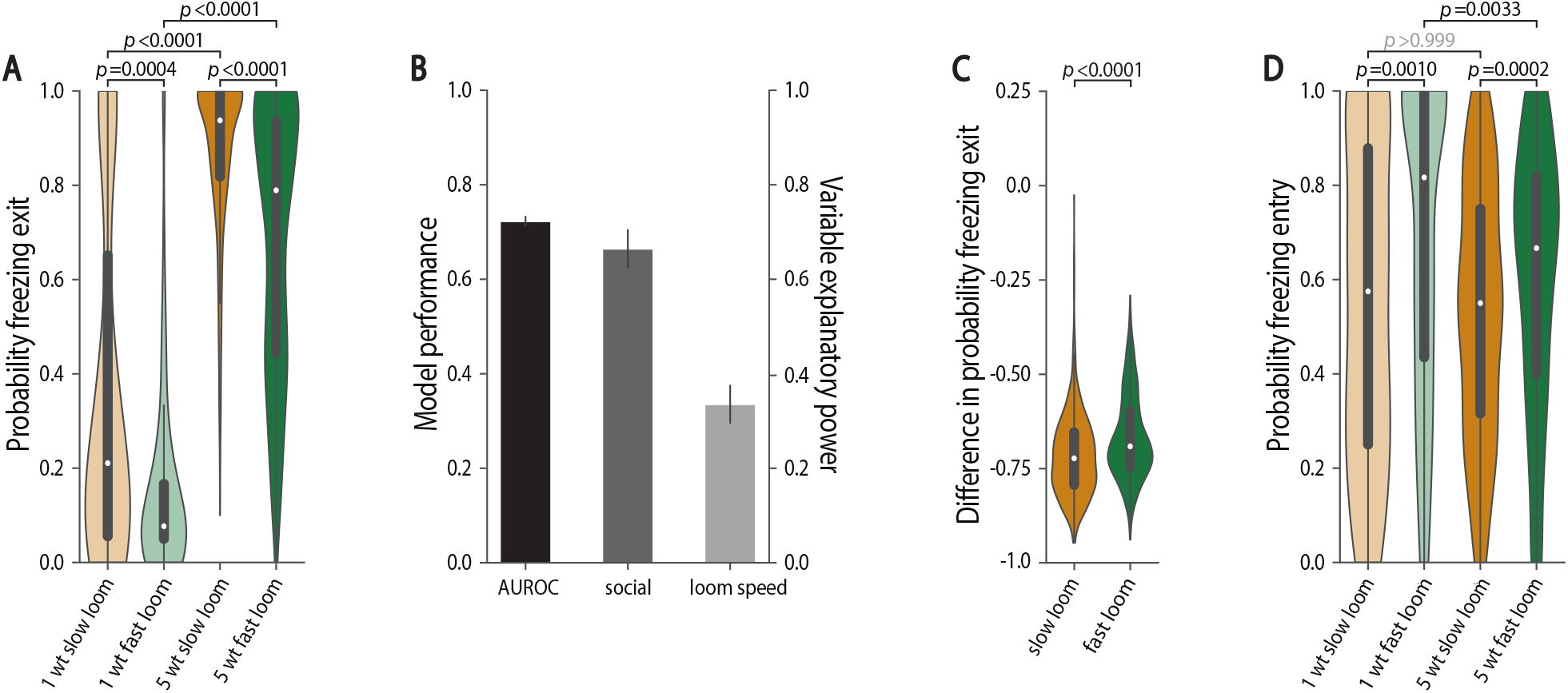
Probability of freezing exit in groups scales with threat imminence. A, C, D) Violin plots representing the probability density distribution of individual fly data bound to the range of possible values, with boxplots elements: center line, median; box limits, upper (75) and lower (25) quartiles; whiskers, 1.5x interquartile range). A) Probability of freezing exit before the following looming stimulus. P-values result from Kruskal–Wallis statistical analysis followed by Dunn’s multiple comparisons test. B) Logistic regression model of the decision to stop or continue freezing as a function of social motion cues and looming speed (10,000 bootstrapping events). Mean and standard deviation of model performance (AUROC – area under the receiver operating characteristic curve, black) and explanatory power of the social environment (dark gray) and looming speed (light gray). C) Difference in the probability of freezing exit between individually tested flies and flies tested in groups for slow and fast looms (see Methods). P-value results from two-tailed Mann–Whitney test. D) Probability of freezing entry upon looming stimulus. P-values result from Kruskal–Wallis statistical analysis followed by Dunn’s multiple comparisons test.

To further explore the impact of threat level on the decision to stop freezing we used a logistic regression model. In our prior study ^21^ such a model revealed that the motion cue generated by surrounding flies was the strongest predictor of the decision to stop freezing, explaining close to 90% of the variance in the data. Therefore, here we modelled freezing exits, using the motion cue of others and looming speed as predictors (see methods, Figure 2B). This model accurately describes our data, as seen by the area under the receiver operating characteristic (AUROC), a measurement of model accuracy (0.72 ± 0.011), and shows that although the social motion cue explains most of the variance in the data (0.66 ± 0.041), looming speed also explains a significant part (0.33 ± 0.041). This model does not however allow us to look at the interaction between looming speed and the impact of social environment on the decision to stop freezing and resume activity. To address this issue, we compared the impact of the social environment on freezing exits observed for flies exposed to fast and slow looms, that is we computed the difference P(Fexit)individual - P(Fexit)social, for flies exposed to both loom speeds, where a negative value means there are more freezing exits in flies tested in groups. We found a small but reliable difference across loom speeds, as seen by the less negative values for flies exposed to the faster rather than to the slower looms (see Methods, MW p<0.0001, Figure 2C). Thus, the social environment had a stronger impact on freezing exits of flies exposed to slow looms, albeit to a small degree, in line with the finding that both for flies tested individually and for flies in groups, freezing bouts are longer when exposed to fast looms than when exposed to slow looms (Supplementary Figure 1C and D). However, it stands in contrast with the stronger social impact on total time spent freezing for flies exposed to the fast loom, suggesting that the probability of freezing upon loom may be decreased in groups to a larger extent during exposure to faster looms. Indeed, we found that flies exposed to slow looms individually or in groups are equally likely to enter freezing, whereas for flies exposed to fast looms the being in a group decreases the probability of freezing entry (p<0.0001, Figure 2D). This finding could result from a shift in balance of the weights given to social danger and safety cues as a function of threat level. In conclusion, looming speed affects both freezing entries and exits, an effect that may interact, even if weakly, with the impact of the social environment on freezing behavior.

Our results so far establish that both threat imminence and the social environment affect freezing responses. However, in these experiments the social environment, i.e. the behavior of flies in the group, varies with threat level, as reflected in the increased average motion cue generated by groups of flies exposed to the slow looms relative to the motion cues produced by groups exposed to fast looms (Supplementary Figure 2A). Therefore, to test the impact of threat imminence on the use of social cues, it is crucial to have experimental control over the social cues, namely the motion of others, such that for different threat levels the motion cues surrounding a focal test fly remain similar. To manipulate the social environment, we controlled the proportion of moving and freezing flies around a focal fly, from all four flies moving to all freezing and the various proportions in between. The same group compositions were exposed to the fast and slow looms (Figure 3). We used blind, NorpA mutant flies, which do not perceive the looming stimulus and walk the entirety of the experimental time, as the moving neighboring flies. This produces the highest surrounding motion cues, a manipulation we had previously shown to lower freezing by focal flies ^21^. To create freezing flies, we artificially induced freezing by optogenetically activating lobula columnar neurons 6 (LC6) using the channelrhodopsin CsChrimson ^44^, LC6>CsChrimson (Supplementary Figure 3A). Making use of these freezing and moving fly lines to create different proportions of moving and freezing flies (Supplementary Figure 3B and C), we were able to produce graded motion cues in groups for both looming speeds (Figure 3A and B). Crucially, these motion cues were similar across threat imminences (Figure 3C). However, optogenetically activating LC6 neurons in addition to driving freezing, also triggers jumps coupled to the loom presentations, especially during the first two seconds of stimulation (Supplementary Figure 3D and E). This means that stable graded motion cues were present about two seconds after the first loom, being briefly interrupted upon loom onset subsequently (Figure 3A and B). It was therefore not possible with this manipulation to examine freezing as a social cue of threat, which mostly modulates freezing onset ^21^. However, the stable motion cues in between looming stimuli allowed the study of the use of safety cues, which modulate the resumption of activity by a fly that froze after the loom ^21^.

**Figure 3.**
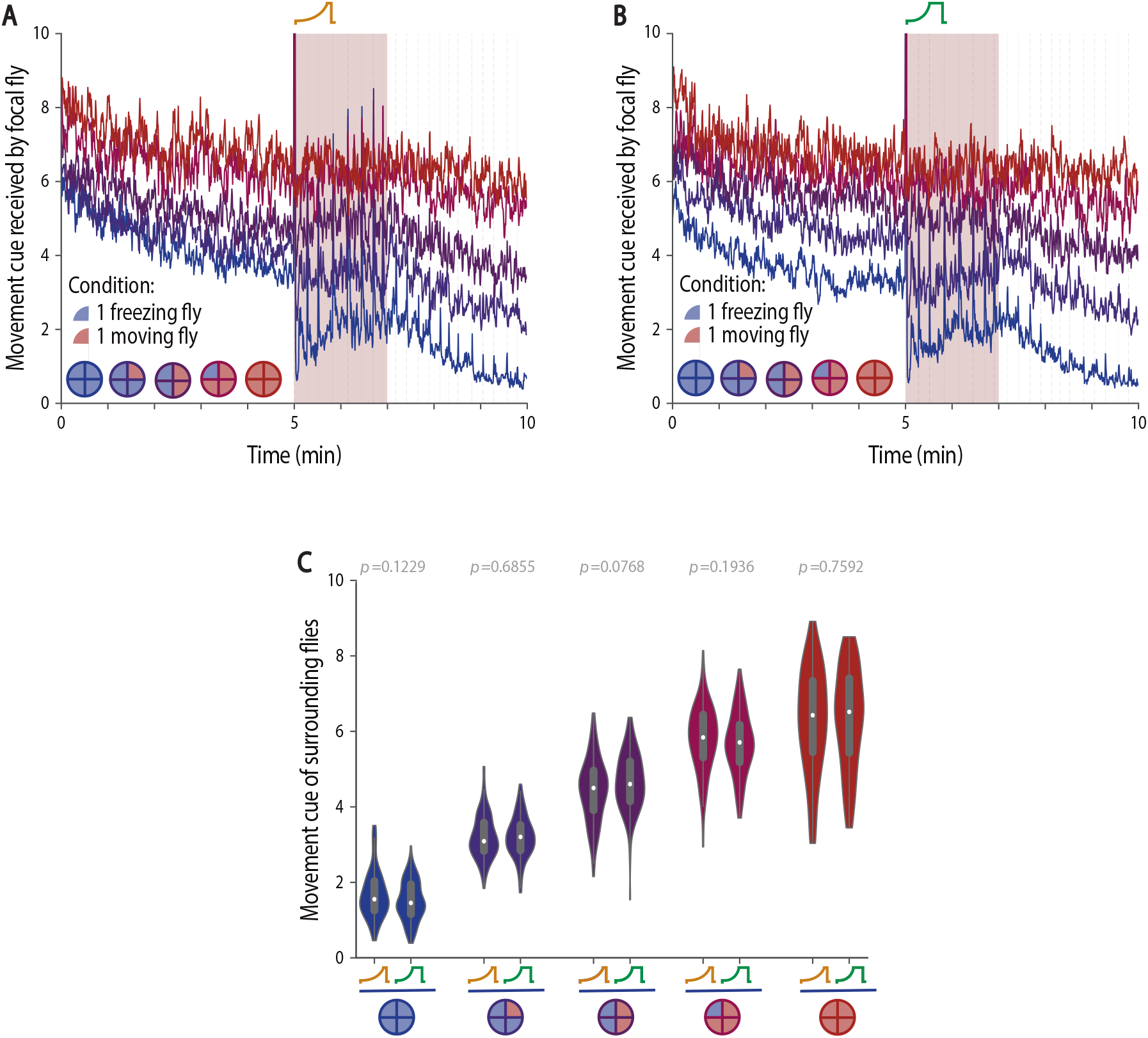
Manipulating the social environment produces similar motion cues across threat imminences. A-C) We manipulated four out of the five flies in group, to surround focal flies with groups with different proportions of flies that always move (blind flies, NorpA, red) and flies that are optogenetically made to freeze (LC6>CsChrimson, blue). The color code for the groups is presented in A and B. A-B) Motion cues produced by the manipulated surrounding flies throughout the experiment when exposed to slow (A) or fast looming stimuli (B); dashed lines represent looming stimuli presentations. C) Violin plot representing the probability density distribution of individual fly data bound to the range of possible values, with boxplots elements: center line, median; box limits, upper (75) and lower (25) quartiles; whiskers, 1.5x interquartile range). Average motion cues produced by the manipulated surrounding flies during the stimulation period. P-values result from two-tailed Mann–Whitney test; significance is determined via Bonferroni correction.

Having a handle on the social environment allowed us to analyze the behavior of focal wildtype flies exposed to similar social motion cues while being presented with looming stimuli of different speeds (Figure 4) and assess how different threat levels affect social cue usage. In both cases we found that a graded manipulation of the social motion cues leads to graded freezing responses of focal flies, as seen in the proportion of flies freezing throughout the experiment (Figure 4A and B) and in the proportion of time each fly spends freezing (KW p < 0.0001; Figure 4C). In addition, overall, flies exposed to faster looms freeze more than flies exposed to slower looms (Figure 4C). In opposition to our previous findings ^21^, flies surrounded by all freezing flies froze less than flies alone; we believe this is due to the induction of jumps, which affects freezing entries as mentioned above.

**Figure 4.**
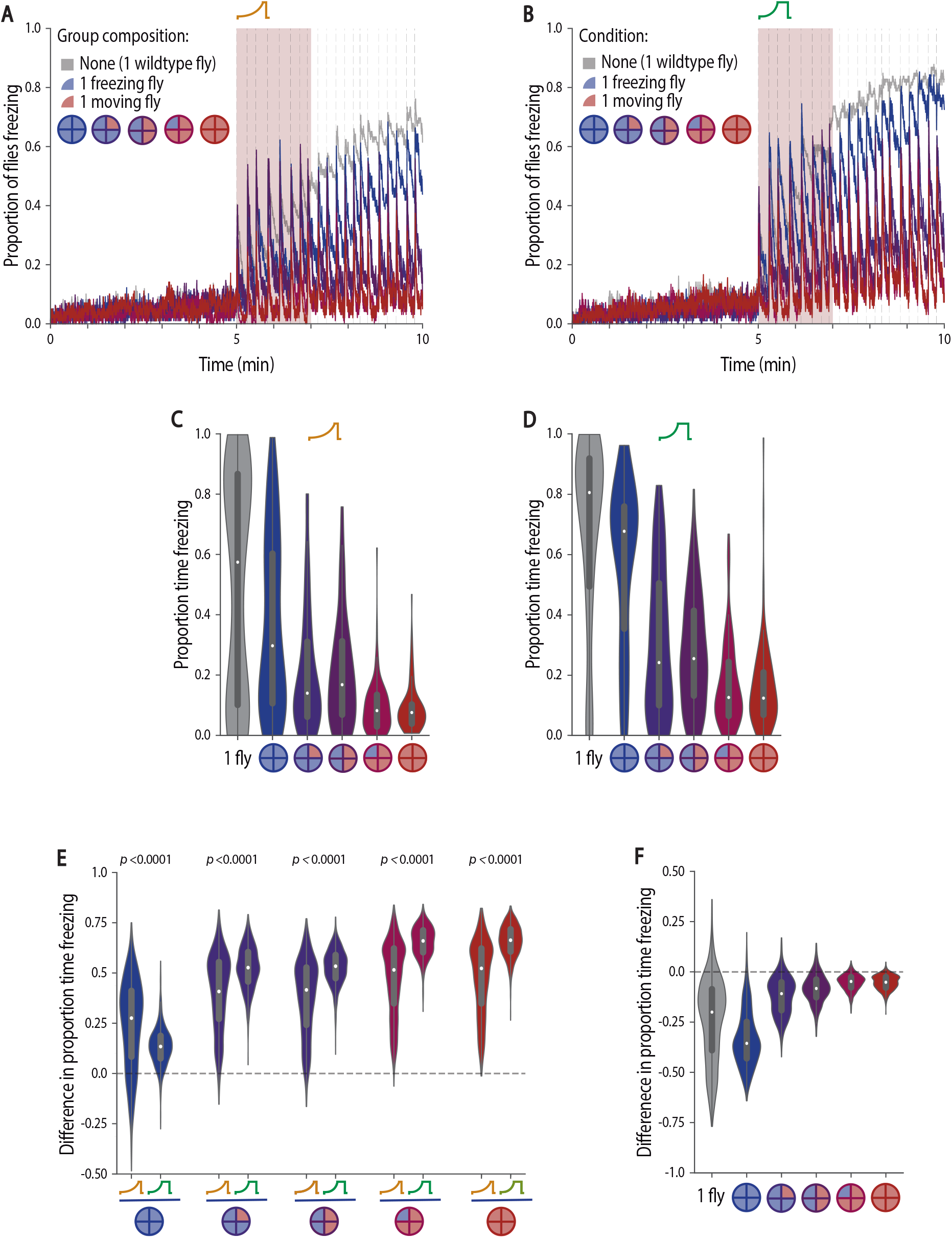
Social environment overrides the perceived imminence of a threat in guiding freezing responses. A-F) We surrounded focal flies with four manipulated flies creating groups with different proportions of flies that always move (blind flies, NorpA, red) and flies that are optogenetically made to freeze (LC6>CsChrimson, blue). The diagrams of circles divided in four depict the numbers of moving and freezing flies, and the outline color represents the colour used to identify group composition; grey shadings represent an individually tested wild-type fly. A-B) Fraction of focal flies freezing throughout the experiment when exposed to slow (A) or fast looming stimuli (B); dashed lines represent looming stimuli presentations. C-F) Violin plots representing the probability density distribution of individual fly data bound to the range of possible values, with boxplots elements: center line, median; box limits, upper (75) and lower (25) quartiles; whiskers, 1.5x interquartile range). C-D) Proportion of time spent freezing in the stimulation period when exposed to slow (C) and fast (D) looms. E) Difference in the proportion of time spent freezing between individually tested flies and flies tested in groups for slow and fast looms (see Methods). P-values result from two-tailed Mann–Whitney test. F) Difference in the proportion of time spent freezing between focal flies exposed to the two looming speeds, for each group composition (see Methods). Statistical comparisons between conditions are presented in Supplementary Table 1.

Importantly, we could now compare the impact of similar social cues across different threat levels, by plotting the difference between freezing by focal flies tested individually and in each social condition, for both fast and slow looms (see Methods; Fig 4E). We found that the impact of the social environment was stronger when flies were exposed to the faster loom in the presence of moving flies that provide social cues of safety, as the differences relative to individually tested flies were bigger (MW p<0.0001; Fig 4E).

As mentioned above, freezing responses scale with threat imminence, but the social environment seems to have a bigger weight in the presence of a higher threat level, which may result from stronger impact of the social safety cues. Indeed, when comparing the difference in freezing responses between looming stimuli, across the graded social motion cues, it is evident that there is an effect of looming speed on the time spent freezing (see Methods; KW p < 0.0001; Figure 4F, Supplementary Table 1), but that this effect decreases and flattens out with the addition of moving flies to the social environment (Prob(Fexit) = 0.1–10.079 IQR −0.19--0.26, Figure 4F, Supplementary Table 1). To summarize, adding moving flies, hence adding motion cues, levels out differences in freezing responses across looming speeds.

To further understand the effect of these tightly controlled social cues on freezing responses in groups we once again focused on freezing exits in between loom presentations (Figure 5). Overall, for both looming speeds, increasingly adding motion cues, by adding moving flies, leads to an increase in the probability of freezing exit (KW p < 0.0001; Figure 5A and B). The stronger impact of social cues of safety on faster looms is again evident comparing the magnitude of the difference between freezing exits by focal flies tested individually and in the presence of moving flies for both fast and slow looms (MW p<0.0001; Figure 5C). The finding that flies exposed to strong motion safety cues show decreased total time spent freezing and increased probability of freezing exits, for both loom speeds, indicates that the length of individual freezing bouts is decreased. Indeed, for all social conditions freezing bouts are shorter the more moving flies are around (Figure 5E and F) and hence the bout length depends on stimulus strength, i.e., the level of surrounding motion cues. This suggests flies gather information about safety for longer, when this information is sparser. Interestingly, at all difficulty levels, the distribution of these freezing bout lengths exhibited positive skew, a characteristic of information accumulation to threshold ^46–49^. In addition, though there are differences in the probability of freezing exit across looming speeds for all group compositions, with slower looms inducing higher freezing exit probabilities, these differences once again become very small as soon as one moving fly is added (Prob(Fexit) = 0.12–0.033 IQR −0.012– 0.17, Figure 5D, Supplementary Table 2). Interestingly, social cues of danger provided by 4 freezing flies seem to be more salient for flies exposed to faster looms (Figure 5C and D); however, an appropriate analysis of this effect warrants a different experimental design in which jumps do not confound the analysis. In conclusion, social cues of safety lead to similar probabilities of freezing exiting across looming speeds and hence override differences in threat level.

**Figure 5.**
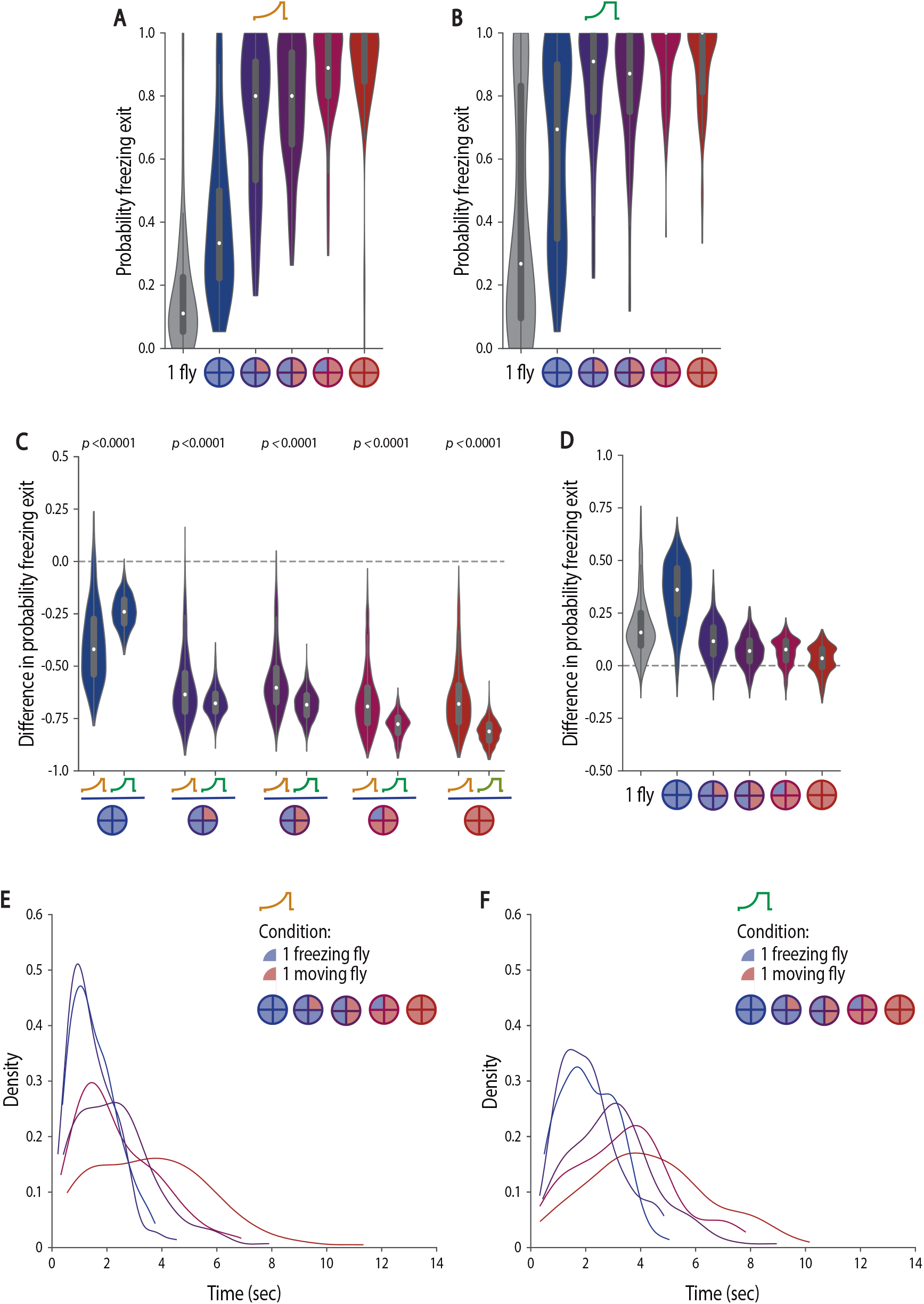
Social cues of safety lead to freezing breaks which underlie freezing response similarities across threat imminences. A-F) We surrounded focal flies with four manipulated flies creating groups with different proportions of flies that always move (blind flies, NorpA, red) and flies that are optogenetically made to freeze (LC6>CsChrimson, blue). The color code for the groups is presented in A and B; grey shadings represent an individually tested wildtype fly. A-D) Violin plots representing the probability density distribution of individual fly data bound to the range of possible values, with boxplots elements: center line, median; box limits, upper (75) and lower (25) quartiles; whiskers, 1.5x interquartile range). A-B) Probability of freezing exit before the following looming stimulus when exposed to slow (A) or fast looming stimuli (B). C) Difference in the probability of freezing exit between individually tested flies and flies tested in groups for slow and fast looms (see Methods). P-values result from two-tailed Mann–Whitney test. D) Difference in the probability of freezing exit between focal flies exposed to the two looming speeds, for each group composition (see Methods). Statistical comparisons between conditions are presented in Supplementary Table 2. E-F) Kernel density estimate plots of the distribution of freezing bout lengths for flies exposed to slow (E) and fast (F) looms.

## Discussion

In this study we have addressed how threat imminence impacts defensive behaviors in groups and reliance on social cues. We show that flies respond with different freezing levels to looming stimuli approaching at different speeds, whether tested individually or in groups, with faster looms triggering faster and more sustained freezing responses. Interestingly, we identified a non-linear scaling effect of looming speed, that is the increase in freezing caused by exposure to a higher threat is not similar across social conditions. This increase in freezing was more pronounced for flies tested individually than flies tested in groups, indicating that groups of flies exposed to faster looms showed a stronger social buffering effect than groups exposed to slower looms. Moreover, controlling the social cues surrounding a fly through the manipulation of group composition under both looming conditions, revealed that social cues of safety override differences in freezing responses to the two threat levels.

With the manipulation of looming speeds, we observed that faster looms lead to a faster engagement in freezing responses, which are then maintained for more prolonged periods of time, whether flies are tested individually or in groups. These findings are consistent with a perceived higher threat level for faster looms leading to a more vigorous response whose intended outcome is undetectability by a potential predator until safety is established. The differences between the latencies to start freezing are in line with differences in latencies to escape in fish ^33^. On the other hand, freezing duration increases with faster looming speeds at an apparent contrast with the reported shorter escape duration for fast looms. However, a closer examination of the flies’ behavior suggests that at a functional level the change is in the direction of increased protection. Flies exposed to an escapable fast loom cut short their sequence of preparatory behaviors that ensures a controlled take-off flight away from the predator, resulting in a faster take-off, albeit less controlled. The small difference in take-off duration may grant flies precious time to survive the chase ^50^. When exposed to an inescapable fast loom flies freeze for a longer period of time, which may allow them to remain undetected in case the predator looms again in a second chase attempt.

Importantly, we uncovered a hitherto unknown non-linear scaling effect of defense responses in groups with threat imminence. The enhanced social buffering effect upon fast looms seems to result from the perception of the approach speed of the threat in interaction with the perception of surrounding social cues, raising the question of whether this effect is generalizable across different features of predation threat level or rather specific to approach speed. To address this issue, it will be interesting to analyze freezing responses tampering with various features of predation threat.

Crucially, we identified that a graded manipulation of the social environment, providing graded levels of motion cues, induces graded freezing responses. Furthermore, underlying the graded amount of total time spent freezing is the modulation of the probability of freezing exit, resulting in graded freezing bout durations. These freezing bout durations uncover progressively faster freezing disengagement, that is, decreasing reaction times to increasing safety motion cues, for both threat imminences. With these findings we uncovered a different strategy to that observed in social copying in the context of feeding and reproduction-related decisions ^17,51,52^, where animals adopt a conformity strategy, following the decision of the majority of others. If flies in our experiments were conforming to the majority one would expect higher freezing levels for groups with 3 and 4 freezing flies, which was not the case. The finding that flies in our experimental conditions do not conform to the group majority and show stronger social buffering when exposed to a higher threat level may seem surprising. It is possible however that very sustained freezing responses, several minutes at a time, may become too costly ^23^ and that responding to the social environment may reduce the cost without significantly reducing the flies’ defenses. The behavioral pattern we observed is consistent with such a strategy, as flies responded to the looming stimulus with freezing even in groups with moving flies; however, each incremental addition of social safety cues lead them to disengage from freezing after increasingly shorter times. This pattern is reminiscent of a process of evidence integration, of safety cues, to decision bound, resumption of activity. A finer grained investigation of how freezing responses of individuals and flies in groups vary with threat level, with careful control and monitoring of motion cues, will permit determining whether freezing responses in flies follow an integrate to threshold model of decision making, analogously to that observed for escapes decisions in mice ^53^ and two-alternative forced choices visual and olfactory tasks in primates, rodents and flies ^46–49^.

Interestingly, social cues of danger produced by surrounding freezing flies seem to exacerbate freezing responses to the faster loom compared with the slower loom since the former are a lot less likely to exit freezing. However, our experiments do not allow addressing the social effect on freezing onset appropriately since the optogenetic manipulations used also produce strong jumping responses. Future experiments inducing freezing without jumps will allow studying the interplay between threat levels and social cues of danger.

We believe this study opens up a path to understand the dynamics of the usage of individually-perceived information and social safety and danger cues in different predation settings, which will provide valuable insight into how crucial threat response decisions are made.

## Acknowledgements

We would like to thank the Fly Platform at the Champalimaud Centre for the Unknown for providing fly work related infrastructure, expertise, and support; the Moita lab for helpful discussions and Anna Hobbiss in particular for pilot experiments with looms of different speeds; João Frazão for expertise and support developing acquisition and tracking codes in Bonsai. This work was supported by Fundação Champalimaud, by Fundação para a Ciência e a Tecnologia (FCT) project UIDB/04443/2020 and ERCCoG819630-A-Fro, as well as by the research infrastructure CONGENTO LISBOA-01-0145-FEDER-022170. Mirjam Heinemans and Matheus Farias were supported by fellowships from FCT, respectively, SFRH/BD/143423/2019 and SFRH/BD/130320/2017.

## Author contributions

C.H.F. and M.A.M. conceived the project and designed the experiments with input from M.H.; C.H.F. and M.H. performed all experiments except for optogenetic activation in the absence of looming which was done by M.F.; R.G. made fly crosses and prepared flies for the experiments. M.F. updated behavioral setups and video acquisition codes. M.F. and M.H. wrote tracking and behavior classification codes. C.H.F., M.H. and M.F. wrote analysis code. C.H.F. and M.H. analyzed the data. C.H.F., M.H. and M.A.M. discussed results. C.H.F. and M.A.M. wrote the manuscript; all authors commented on the manuscript.

## Inclusion and diversity

One or more of the authors of this paper self-identifies as a member of the LGBTQ+ community.

