## Supplemental Figure1 for "Social cues of safety can override differences in threat level"

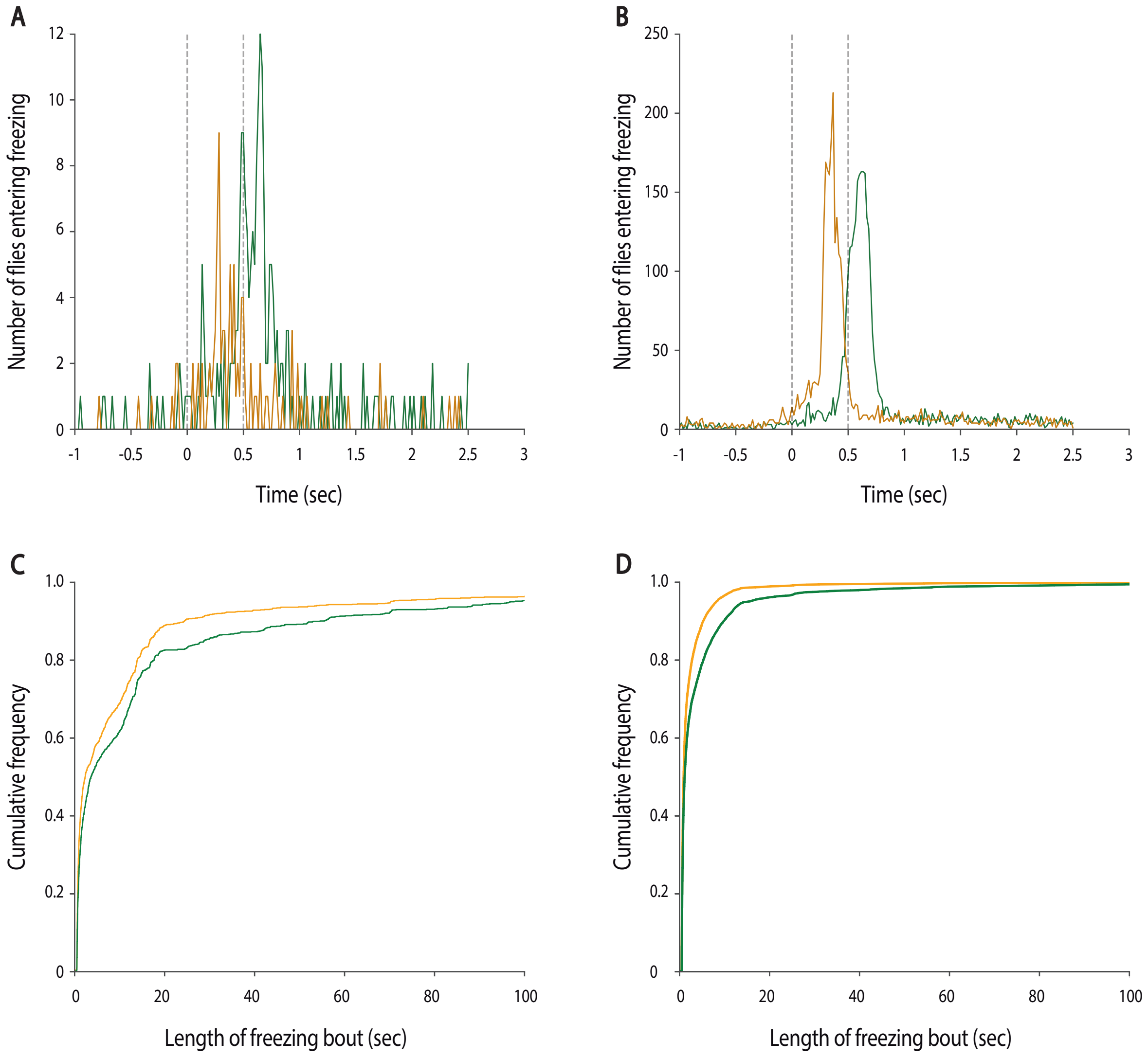


Supplementary Figure 1 – **Effect of looming speed on latency to freezing onset and freezing bout length.** A-B) Distribution of the timepoints of freezing onset after looming for flies tested individually (A) and in groups (B). C-D) Cumulative distributions of freezing bout lengths for flies tested individually (C) and in groups (D). Orange denotes flies exposed to slow looms and green flies exposed to fast looms.
