## Supplemental Figure2 for "Social cues of safety can override differences in threat level"

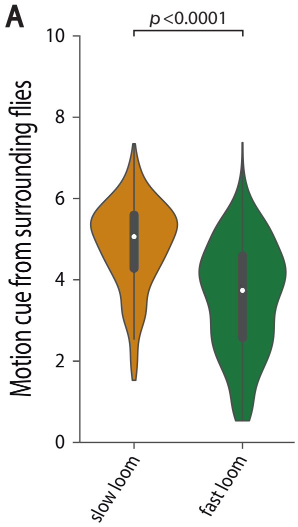


Supplementary Figure 2 – **Motion cues in groups scale with threat intensity.** A) Violin plots representing the probability density distribution of individual fly data bound to the range of possible values, with boxplots elements: center line, median; box limits, upper (75) and lower (25) quartiles; whiskers, 1.5x interquartile range). Average motion cue a focal fly is exposed to during the stimulation period. P-value results from two-tailed Mann–Whitney test.
