## Supplemental Figure3 for "Social cues of safety can override differences in threat level"

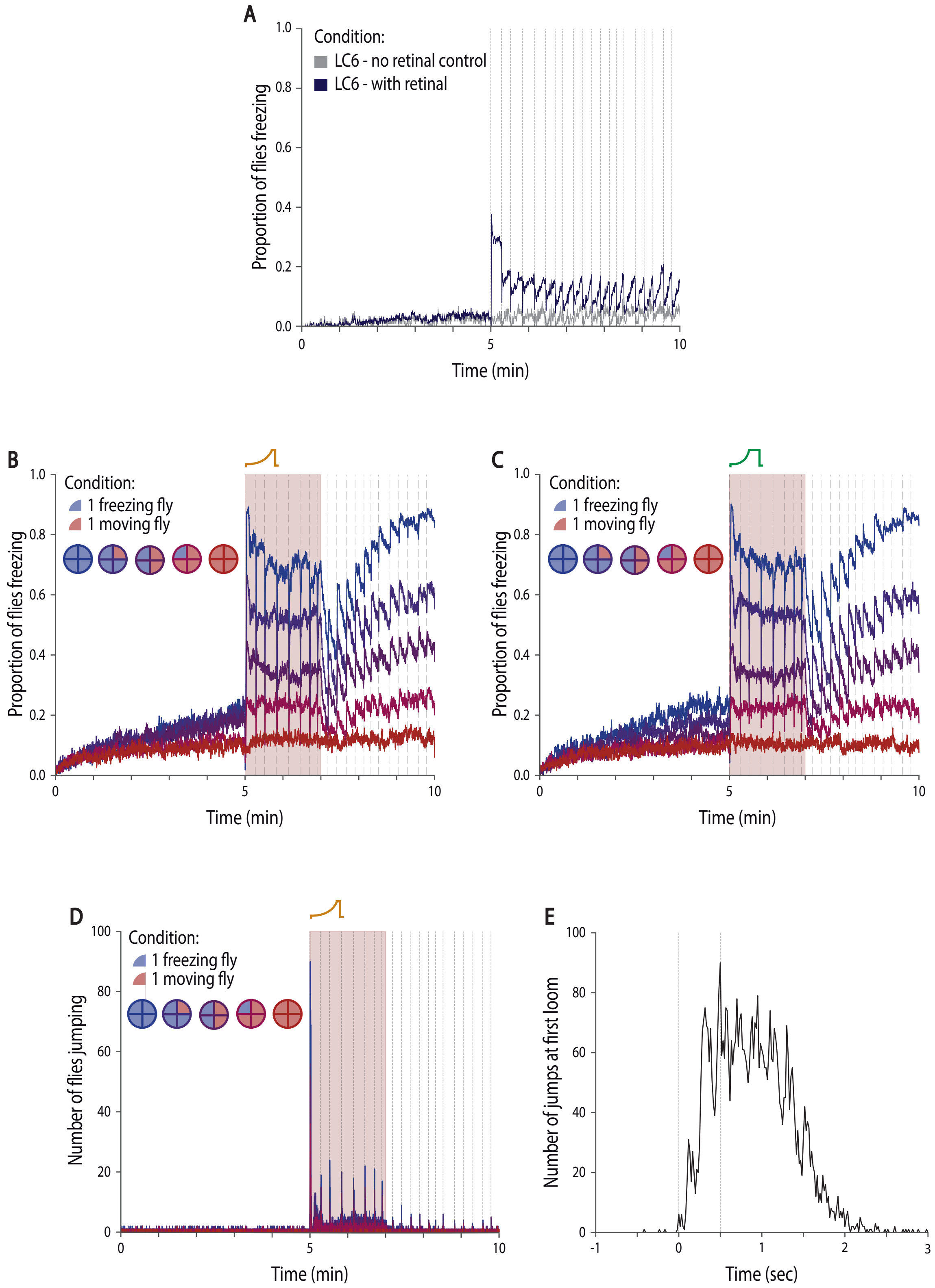


Supplementary Figure 3 – **Freezing and jumping responses of optogenetically activated LC6>CsChrimson.** A) Fraction of flies freezing throughout the experiment while providing pulsed red light at the timestamps normally used to provide looming stimuli (dashed lines); LC6>CsChrimson flies supplemented with retinal (blue) and control without (grey). B-D) We manipulated four out of the five flies in group, to surround focal flies with groups with different proportions of flies that always move (blind flies, NorpA, red) and flies that are optogenetically made to freeze (LC6>CsChrimson, blue), while presenting looming stimuli. The color code for the groups is presented in B, C and D. B-C) Fraction of surrounding, manipulated, flies freezing throughout the experiment when exposed to slow (B) or fast looming stimuli (C); dashed lines represent looming stimuli presentations. D-E) Data from flies exposed to slow looms. D) Number of jumps throughout the experiment by the surrounding manipulated flies. E) Number of jumps at the first loom presentation for LC6>CsChrimson flies in groups of four surrounding optogenetically manipulated flies.
