## Supplemental Table1 for "Social cues of safety can override differences in threat level"

Table 1. Statistical comparisons of the difference in time each fly spent freezing when exposed to 25 and 50 cm/s looming stimuli, tested individually and in groups of freezing and moving flies (p-values for Kruskall-Wallis followed by Dunn's multiple comparisons test).

| | 4 freezing | 3 freezing | 2 freezing | 1 freezing | 0 freezing |
|---|---|---|---|---|---|
| 1 fly alone | <0.0001 | <0.0001 | <0.0001 | <0.0001 | <0.0001 |
| 4 freezing | | <0.0001 | <0.0001 | <0.0001 | <0.0001 |
| 3 freezing | | | <0.0001 | <0.0001 | <0.0001 |
| 2 freezing | | | | <0.0001 | <0.0001 |
| 1 freezing | | | | | >0.9999 |
