## Supplemental Table2 for "Social cues of safety can override differences in threat level"

Table 2. Statistical comparisons of the difference in the probability each fly has of exiting freezing when exposed to 25 and 50 cm/s looming stimuli, tested individually and in groups of freezing and moving flies (p-values for Kruskall-Wallis followed by Dunn's multiple comparisons test).

| | 4 freezing | 3 freezing | 2 freezing | 1 freezing | 0 freezing |
|---|---|---|---|---|---|
| 1 fly alone | <0.0001 | <0.0001 | <0.0001 | <0.0001 | <0.0001 |
| 4 freezing | | <0.0001 | <0.0001 | <0.0001 | <0.0001 |
| 3 freezing | | | <0.0001 | <0.0001 | <0.0001 |
| 2 freezing | | | | >0.9999 | <0.0001 |
| 1 freezing | | | | | <0.0001 |
